# Mutation of phosphatidate phosphohydrolase genes confers broad-spectrum disease resistance in plants

**DOI:** 10.1101/2024.07.04.602136

**Authors:** Qiuwen Gong, Gan Sha, Xinyu Han, Zhenhua Guo, Lei Yang, Wei Yang, Ronglei Tan, Guang Chen, Yufei Li, Xin Shen, Kabin Xie, Guangqin Cai, Honghong Hu, Jie Luo, Qiang Li, Guotian Li

## Abstract

Phosphatidic acid (PA) is considered as a second messenger that interacts with protein kinases, phosphatases and NADPH oxidases, amplifying the signal to initiate plant defense signaling responses (Li and Wang, 2019). In rice, mutation of *RBL1* causes the accumulation of PA, enhancing multipathogen resistance (Sha et al., 2023). In our previous study, we attempted to rescue *rbl1* mutant by overexpressing phosphatidate phosphohydrolase (*PAH*) genes. However, overexpression of *PAH2* reduced the PA level but did not affect the disease resistance, which made us to reconsider the importance of PA and *PAH* in rice immunity. Here, we identified that mutation of *PAHs* caused PA accumulation and enhanced multipathogen resistance in rice and *Arabidopsis*.

---

Phylogenetic analyses reveal that PAHs are highly conserved in plants. Rice PAHs contain conserved NLIP and LNS2 signature motifs (Figure 1a). The gene expression assays showed that *PAH1* and *PAH2* were transcribed in all tissues examined, with the highest levels in the leaf (Figure 1b). Subcellular localization assays indicated that PAH-GFP signals predominantly co-localized with the endoplasmic reticulum (ER) marker HDEL1 (Figure 1c).

**Figure 1.**
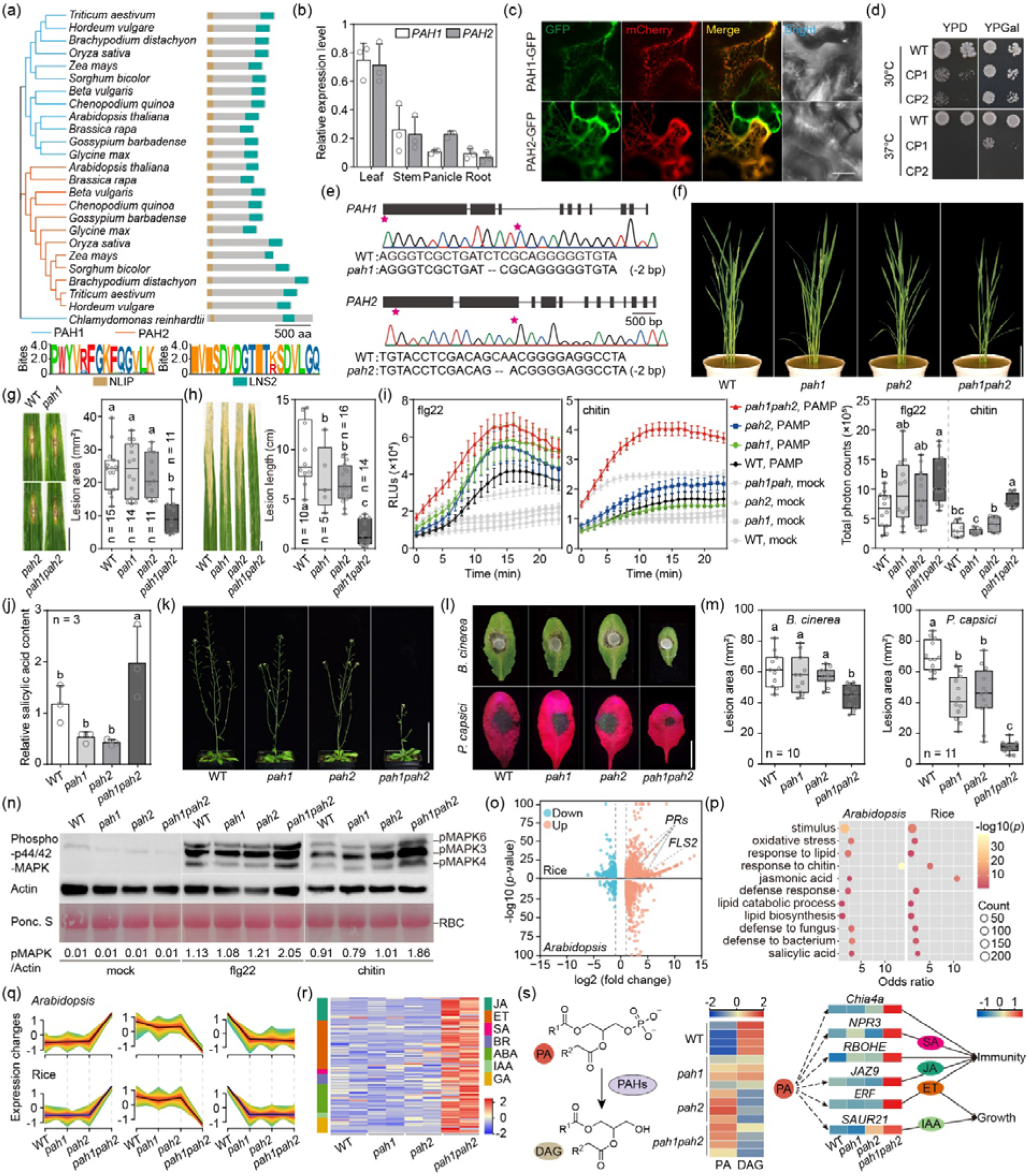
Mutation of the *PAH* genes confers broad-spectrum disease resistance in both rice and *Arabidopsis*. (a) Phylogenetic analyses of PAH homologs from plant species. The NLIP and LNS2 motifs are shown at the bottom. (b) qRT-PCR assays of *PAH1* and *PAH2*. (c) Subcellular localization assays of rice PAHs. Bar = 10 μm. (d) Expression of rice *PAH1* rescues the growth defect of yeast *pah1* mutant. (e) Structures and gene-editing sites (asterisks) in rice *PAH1* and *PAH2*. (f) 9-week-old WT, *pah1, pah2*, and *pah1pah2* rice plants. Bar = 15 cm. (g-h) Lesions caused by *Magnaporthe oryzae* (g) and *Xoo* (h) at 14 days post-inoculation. Bars = 1 cm. (i) ROS production in the *pah1, pah2, pah1pah2*, and WT plants. PAMPs are flg22 or chitin as indicated. (j) The total SA level is increased in the *pah1pah2* mutant. (k) The *Arabidopsis pah* mutants and Clo-0 plants at the flowering stage. Bar = 10 cm. (l-m) Infected leaves (l) and lesion areas (m) of WT and *pah* lines inoculated with pathogens. Bars = 1 cm. (n) Immunoblotting analyses of MAPK activation. Ponceau S staining indicates the protein loading. (o) Volcano plots of DEGs in rice and *Arabidopsis pah1pah2* mutants. (p) GO enrichment analyses of DEGs in (o). (q) Hierarchical clustering analyses of genes in *Arabidopsis* and rice *pah* lines. (r) Heatmap of the expression levels of hormone-related genes in *pah1pah2*. (s) Roles of the PAHs in PA metabolism, rice growth and immunity. The heatmap indicates the PA and DAG contents, and four biologically independent repeats for each line. These color blocks beneath each gene indicate the gene expression level. Different letters were calculated using the Duncan’s new multiple range test. n = number of biologically independent repeats.

In yeast, the *pah1* mutant is lethal at 37°C. We transformed rice *PAH1* and *PAH2* genes into the wild-type (WT) yeast strain, respectively, and knocked out the yeast endogenous *PAH1* gene. The rice *PAH1* complementation strain (CP1) grew well on the induction medium YPGal but not on the non-inducing medium YPD at 37°C, indicating that rice *PAH1* functions as a PA phosphohydrolase in yeast. In contrast, rice *PAH2* gene was unable to complement the yeast *pah1* mutant (Figure 1d).

To investigate the functions of rice *PAHs*, we genome edited *PAHs* by targeting their first exons (Figure 1e). We then crossed the homozygous *pah1* and *pah2* lines and obtained the *pah1pah2* double mutant plants. The plant height of *pah1pah2* was reduced to 78% of the WT (Figure 1f, Figure S1). We next tested the disease resistance of rice *pah* with rice blast pathogen *Magnaporthe oryzae*. The infected area of *pah1pah2* was only 58% of the WT (Figure 1g). Similarly, the *Xanthomonas oryzae* pv. *oryzae* (*Xoo*) infection assays showed that the lesions were much shorter in *pah1pah2* (1.15 cm) than the WT (9.40 cm) (Figure 1h). In summary, both fungal and bacterial infection assays demonstrate enhanced resistance of rice *pah1pah2*. Subsequently, we used flg22 and chitin to investigate the ROS production in *pah* plants. The results showed that ROS production in *pah1pah2* line was 1.5- and 3.5-fold more than that in WT when challenged with flg22 and chitin, respectively, and the total photon counts, also denoting the ROS level, of *pah1pah2* line showed 1.78- and 2.58-fold increases (Figure 1i). Salicylic acid (SA) plays a key role in plant defense. The quantification assays showed that the SA level increased 1.68-fold in *pah1pah2* compared with the WT (Figure 1j).

To investigate whether the role of the *PAH* genes in immunity is conserved, we obtained *Arabidopsis pah* T-DNA insertion mutants (Eastmond et al., 2010). At 5 weeks post-sowing, the *pah1pah2* seedlings were shorter than the WT (Col-0) (Figure 1k). Infection with *Botrytis cinerea, pah1pah2* plants developed smaller lesion areas, a 28.3% reduction than the WT. Similarly, when inoculated with *Phytophthora capsici*, the lesion area of *pah1pah2* (11 mm^2^) was only 15.7% of the WT (70 mm^2^) (Figure 1l, m). We further examined the levels of phosphorylated MAPKs (pMAPKs). Under flg22 and chitin treatments, the levels of pMAPKs were significantly higher in the *Arabidopsis pah1pah2* than that in the WT (Figure 1n). The results are consistent with the increased ROS levels in rice *pah1pah2* mutants.

To investigate *PAH*-mediated regulation of gene transcription and metabolism, we performed RNA-seq and lipidomics analyses. The volcano plots showed that certain *PR* genes and *FLS2* were upregulated in rice *pah1pah2* (Figure 1o). Furthermore, Gene Ontology (GO) enrichment analyses of differentially expressed genes (DEGs) between WT and *pah1pah2* plants showed that ‘response to lipid’, ‘defense response’ and ‘defense response to fungus/bacterium’ were enriched (Figure 1p), which are consistent with the enhanced disease resistance of *pah1pah2* plants. Then using the hierarchical clustering analyses, three clusters were identified in *Arabidopsis* and rice (Figure 1q). In the ‘genes upregulated in *pah1pah2’* cluster, the expression levels of many hormone-related genes that are involved in JA, ET, SA, and IAA, were simultaneously upregulated in *pah1pah2* lines (Figure 1r), indicating that these plant hormones are likely to be involved in resistance and growth of *pah1pah2* lines.

Lipidomics analyses showed that mutation of *PAHs* resulted in the increase of PA and decrease of DAG in rice *pah1pah2* (Figure 1s). Additionally, immunity-related genes, *Chia4a, NPR3, RBOHE, JAZ9* and *ERF*, and genes negatively regulating plant growth, including *SAUR* were upregulated in *pah1pah2* mutants (Figure 1s). Overexpression of the *SAUR* genes inhibited the biosynthesis of plant growth hormones (Xu et al., 2017), some of which were upregulated in *pah1pah2*, thus partially explaining the growth defects of *pah1pah2*. In summary, mutation of *PAH* genes alters phospholipid metabolism in plants, and accumulated PA activates the expression of immunity-related genes, but negatively regulates plant growth.

In conclusion, mutations of both *PAH* genes enhance plant resistance, however, inhibiting plant growth, which shows an immunity-growth tradeoff. Previously, we used genome-editing to break this tradeoff in *RBL1* (Sha et al., 2023), demonstrating that it is feasible to utilize multiplexed genome-editing to generate alleles that balance growth and immunity. Moreover, the use of genome-editing to insert uORFs in the promoter to manipulate protein translation and pathogen-induced silencing of *PAHs* are optional approaches to engineer plants for disease resistance without yield penalty (Sha and Li, 2023; Xue et al., 2023). The PA metabolism genes *PAHs* are highly conserved in plants, and the role of *PAHs* in disease resistance in other crops is worthy of further investigation.

## Acknowledgments

We thank Professor L. Dunkle for the editing of the manuscript. This work was supported by STI2030-Major Projects (2023ZD04070), the Key R&D Program of Hubei Province (2023BBB171), National Natural Science Foundation of China (32172373), Fundamental Research Funds for the Central Universities (2662023PY006 and AML2023A05) and the Open Research Fund of the State Key Laboratory of Hybrid Rice (Wuhan University) (KF202202) to G.L. G.S. is supported by the Open Research Fund of the Guangdong Province Key Laboratory of Microbial Signals and Disease Control (MSDC2023-02). This work was also supported by Hubei Hongshan Laboratory.

## Notes

**Conflict of interest statement** The authors declare no conflict of interest.

### Competing Interest Statement

G.L., G.S., Q.G., and X.H. are coinventors on a patent no. ZL202311556377.0. The remaining authors declare no conflict of interest.

