## Supporting information for "Mutation of phosphatidate phosphohydrolase genes confers broad-spectrum disease resistance in plants"

Additional supporting information may be found online in the Supporting Information section at the end of the article.

Figure S1 Supplementary Figure.

Table S1 Primers used in this study.

**Supporting information**

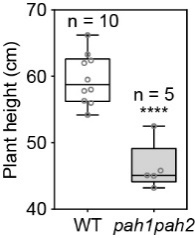

**Supplementary Fig. 1 The rice *pah1pah2* plants display growth defects.**

Plant height of rice wild-type (WT, Kitaake) and *pah1pah2* plants at the flowering stage. Data are displayed as box and whisker plots with individual data points: center line, median; box limits, 25th and 75th percentiles. Asterisks indicate significant differences using the unpaired Student’s *t*-test (*****P* < 0.0001). n = number of biologically independent repeats.

**Supplementary Table S1 Primers used in this study.**

| **Primer** | **Sequence (5’-3’)** |
| --- | --- |
| S3AD5-R | TAGGTCTCCAAACGGATGAGCGACAGCAAAC |
| S5AD5-F | CGGGTCTCAGGCAGGATGGGCAGTCTGGGCA |
| L3AD5-R | TAGGTCTCCAAACGGATGAGCGACAGCAAACAAAAAAAAAAGCACCGACTCG |
| L5AD5-F | CGGGTCTCAGGCAGGATGGGCAGTCTGGGCAACAAAGCACCAGTGG |
| KOPAH1gR2-F | TA GGTCTCC GCTGATCTCGCA GTTTTAGAGCTAGAA |
| KOPAH1gR2-R | CG GGTCTCA CAGCGACCCTAC TGCACCAGCCGGG |
| KOPAH2gR2-F | TA GGTCTCC TCGACAGCAACG GTTTTAGAGCTAGAA |
| KOPAH2gR2-R | CG GGTCTCA TCGAGGTACATGTGCACCAGCCGGG |
| KOPAH1/seqF | CATCCATTCCCCTCCCGTAT |
| KOPAH1/seqR | GTTCAGGCTCACAGCTAGCT |
| KOPAH2/seqF | GGAAGTTCGGGAGCTTCATCT |
| KOPAH2/seqR | CAAGGAGCTCAGCAGCAATC |
| qRT-Actin-F | CAGGCCGTCCTCTCTCTGTA |
| qRT-Actin-R | AAGGATAGCATGGGGGAGAG |
| qRT-PAH1-F | GTCGACGTGAAGTCCTATAC |
| qRT-PAH1-R | CTGCTCAACAAGAGTGGTTG |
| qRT-PAH2-F | GGTGAAGTTGCGGTGAAT |
| qRT-PAH2-R | AGAGCATGGAGGGATGTG |
| PAH1 cDNA-F | ATGAACGTGGTTGGGCGG |
| PAH1 cDNA-R | CAGATCAACATCTGGCAA |
| PAH2 cDNA-F | ATGTACGCGGTGGGG |
| PAH2 cDNA-R | AATATCAACAGCAGGTAACGG |
